# Hyperglycemia and hyperfibrinogenemia alter *Staphylococcus aureus* abscess community morphology, antimicrobial susceptibility, and virulence *in vitro*

**DOI:** 10.64898/2026.02.25.707959

**Authors:** Emily A. Britt, Lyvia K. Markle, Tina I. Bui, Ann Lindley Gill, Benjamin F. Ricciardi, Steven R. Gill

## Abstract

*Staphylococcus aureus* is a prevalent human pathogen responsible for an array of invasive infections, such as osteomyelitis and bacteremia, which may be life threatening, recurrent, and cause permanent tissue damage. *S. aureus* infections are exacerbated in patients with co-morbidities including obesity and type 2 diabetes (obesity/T2D) due to impaired immune function, which leads to chronic inflammation and poor healing. Immune dysfunction can be attributed to gut microbiome dysbiosis, characterized by altered community composition and abundance, with aberrant production of gut-immune axis metabolites. Alongside the heightened infection susceptibility exhibited by obese/T2D hosts, *S. aureus* adapts to the hyperglycemic and hyperfibrinogenemic host environment for robust colonization. *S. aureus* can persist in host tissues by forming staphylococcal abscess communities (SACs) encapsulated by a fibrin pseudocapsule that protect the bacteria from antimicrobials and immune cell killing. Our current work aims to investigate how *S. aureus* adapts to the hyperglycemic and hyperfibrinogenemic obese/T2D environment. Our data show that two *S. aureus* clinical isolates, USA300 FPR3757 and JAR06.01.31, utilize fibrinogen differently in obese/T2D-like conditions to form unique pseudocapsule structures. Furthermore, RNAseq data show that in obese/T2D-like conditions, *S. aureus* upregulates virulence and tissue invasion gene expression. Additionally, our data suggest that antibiotic susceptibility in obese/T2D-like conditions is affected by antibiotic size, charge and metabolic activity of *S. aureus.* Collectively, these investigations will elucidate the impact of hyperglycemia and hyperfibrinogenemia on *S. aureus* abscess formation in two clinically relevant strains and may inform future therapies for obese/T2D patients.

**Importance:** Type 2 diabetes associated with obesity creates a unique host environment that promotes the severity and persistence of *Staphylococcus aureus* infections. Elevated blood glucose and fibrinogen disrupt the normal immune response and create conditions that favor bacterial persistence and dissemination. *S. aureus* is an opportunistic pathogen capable of causing a broad spectrum of diseases ranging from skin infection to life threatening blood and bone infection. A critical step in its pathogenesis is the formation of abscesses, which shield the bacteria from immune clearance and antibiotic treatment. In this study, we demonstrate that the altered metabolic and inflammatory state of the obese diabetic host reshapes the way *Staphylococcus aureus* constructs these protective abscesses. We show that *S. aureus* modifies its use of host fibrin and adjusts its gene expression in response to high blood glucose and fibrinogen, thereby enhancing its ability to persist in host tissues.

## Introduction

Diabetes is a chronic metabolic disease characterized by the inability of the body to produce (Type 1 Diabetes—T1D) or effectively use insulin (Type 2 Diabetes—T2D). Impaired insulin signaling cascades lead to hyperglycemia with decreased innate immune cell function and an altered cytokine response that contributes to a hyperinflammatory environment and a greater incidence of bacterial infections. T2D accounts for 90% of diabetes cases and is associated with socio-economic factors, a sedentary lifestyle, and rising levels of obesity. Diabetics typically have elevated plasma fibrinogen (hyperfibrinogenemia) and reduced fibrinolysis^1–7^. Fifty-one million Americans are living with diabetes, with the population projected to increase to 63 million individuals by 2045^8^.

The hyperglycemic and hyperfibrinogenemic environment of obesity-related type 2 diabetes (obesity/T2D) contributes to a greater risk of invasive and chronic infections of *Staphylococcus aureus*, a bacterial pathogen with a clinical spectrum ranging from asymptomatic skin colonization to osteoarticular infections, endocarditis, and death^2–5,7^. Levels of fibrinogen, an extracellular matrix protein essential for blood clotting^9^, are higher in obese individuals with T2D (from 5.3-7.9 mg/mL^10^) compared to those without T2D^11,12^ (1.5-4.0 mg/mL^13^). Hyperglycemic A1C (Hemoglobin A1C %) levels in obese/T2D patients are often greater than 6.5^14^. Elevated levels of plasma fibrinogen can lead to a hypercoagulable state, increasing the risk of blood clots^13^. *S. aureus* binding of host fibrinogen is an early step in the formation of protective abscesses which enhance survival of the pathogen within the host and dissemination to other organ systems. How *S. aureus* adapts to the obesity/T2D environment and the contribution of increased fibrin deposition on abscess formation remains to be determined.

*S. aureus* abscess formation is a developmental process that begins within hours of infection. Dissemination to organ tissue and release of bacterial lipoproteins initiates Toll-like receptor 2 (TLR2)-mediated inflammatory signals that recruit neutrophils and other immune cells to the infection site. Replication of *S. aureus* within the infected tissue leads to formation of a staphylococcal abscess community (SAC) encapsulated by two concentric structures, an inner fibrin pseudocapsule and an outer fibrin microcolony-associated meshwork (MAM), that function as a protective barrier to peripheral neutrophils^15^. Formation of a SAC is initiated by two extracellular *S. aureus* proteins, coagulase (Coa) and von Willebrand-factor binding protein (vWbp), which activate host prothrombin, forming a thrombin complex that cleaves host fibrinogen into insoluble fibrin^15–18^. Coagulase remains localized to the bacterial surface, where it contributes to formation of the fibrin pseudocapsule. vWbp diffuses away from the bacteria where it forms the MAM^18^. Previous research has demonstrated an adaptive functional response of *S. aureus* to the obese/T2D host^5^. However, modulation of the *S. aureus* transcriptional response and the virulence and metabolic mechanisms associated with SAC formation in the obese/T2D environment are unknown.

To investigate adaptation of *S. aureus* to the obese/T2D host, we propagated *S. aureus* abscesses in an *in vitro* 3D model to assess SAC formation and pseudocapsule structure under defined conditions that mimic obesity/T2D. We hypothesize that hyperglycemia and hyperfibrinogenemia synergistically modulate staphylococcal virulence and SAC architecture. We found that two *S. aureus* isolates, USA300 FPR3757^19^, a soft tissue infection isolate, and JAR06.01.31^20^, a periprosthetic joint infection isolate, uniquely adapt to a hyperglycemic and hyperfibrinogenemic environment by differentially recruiting and utilizing host fibrin to form distinct pseudocapsule architectures. In addition, we found that hyperglycemia and hyperfibrinogenemia drive virulence expression of tissue invasion and immune evasion factors. Finally, we determined that the pseudocapsule is protective of some, but not all, antibiotic challenges. Together, this work demonstrates *S. aureus* adaptation to the obese/T2D environment and provides insights into patient specific treatment strategies.

## Methods

### Animals

All handling of mice and associated experimental procedures were reviewed and approved by the University Committee on Animal Resources at the University of Rochester Medical Center. Male C57BL/6J mice (JAX stock #000664) were housed in five per micro-isolator cages in a two-way housing room on a 12-hour light/dark schedule. At six weeks of age, mice were provided unrestricted access to either lean (10% kcal fat, OpenSource Diets D12450J) or high-fat (60% kcal fat, OpenSource Diets, D12492) diets for 14 weeks, leading to obesity and diet-induced type 2 diabetes for high-fat fed mice^21,22^.

### Human plasma collection

All work with human samples and the protocol for specimen procurement were reviewed and approved by the Research Subjects Review Board at the University of Rochester (IRB STUDY00006496: Type 2 diabetes and *S. aureus*). Samples were collected during preoperative screening of patients undergoing orthopedic surgeries in the Department of Orthopedics at Highland Hospital, University of Rochester Medical Center. In this study, samples with HbA1c between 5.3% - 5.7% were considered healthy and samples with HbA1C > 7% were considered diabetic.

### Bacterial strains and growth conditions

*Staphylococcus aureus* strains USA300 FPR3757^19^ and JAR06.01.31 (number 890; the Culture Collection of Switzerland (CCOS), Wädenswill, Switzerland)^20^ stocks were stored in tryptic soy broth (TSB) containing 20% glycerol at -80°C. Staphylococci were routinely cultured on tryptic soy agar (TSA) and in TSB overnight (18 hours) at 37°C with agitation.

### *S. aureus* growth curves in glucose

Overnight cultures were washed twice in 1X phosphate-buffered saline (PBS) and resuspended to an optical density at 600nm (OD_600_) of 0.5-0.6. The normalized bacteria were further diluted 1:200 in TSB with 0, 5-, 10-, 15-, or 25-mM glucose, in a round-bottom 96-well plate, to a total volume of 200 μL. OD_600_ kinetic readings were measured every 30 minutes with a BioTek ELx808 microplate reader (Agilent Technologies) set at 37°C with medium shaking.

### *S. aureus*-induced clotting assay

Overnight cultures were diluted 1:100 in TSB and grown to stationary phase (OD_600_ ∼1.0). Subcultures were washed twice with PBS and resuspended to approximately 1.0 × 10^8^ CFU/mL. 50 µL of normalized cultures were added to 250 µL of human or mouse plasma in 1.5 mL microcentrifuge tubes and incubated for 24 hours statically at 37°C. After incubation, the clots were scooped out of the microcentrifuge tubes and transferred to sterile, pre-weighted Lysing Matrix A (MP Biomedicals, 116910500) tubes containing PBS to measure clot weights. For CFU enumeration, clots were homogenized with a FastPrep-24 (MP Biomedicals) at 6.0 m/s for 40 s, serially diluted, and plated onto TSA.

### Fibrinogen ELISAs

Fibrinogen concentrations in human and mouse plasma and serum were measured using host-specific enzyme-linked immunosorbent assays (Innovative Research, Mouse #IMSFBGKTT and Human #IHUFBGKT). Plates were precoated with affinity-purified capture antibodies. After washing, biotin-labeled polyclonal anti-human fibrinogen primary antibodies were bound to the captured protein. Excess antibody was washed away with washing buffer and streptavidin conjugated to horseradish peroxidase was added. Tetramethylbenzidine (TMB) substrate was used for color development at 450nm. A standard curve with mouse fibrinogen or human fibrinogen was performed alongside unknown samples.

### *In vitro* collagen model of staphylococcal abscess community (SAC)

Overnight cultures were washed thrice with sterile PBS and resuspended to an OD_600_ of 0.9 ± 0.1 (Beckman, DU530 UV-Vis Spectrophotometer). These cultures were diluted 1:100,000 and mixed into liquid collagen solution. Briefly, rat tail type 1 collagen (RatCol, Advanced BioMatrix #5153) was diluted to a final concentration of 1.78 mg/mL according to manufacturer instructions with provided neutralization buffer, PBS, and diluted culture (∼2.0 x 10^3^ CFU/mL)^18^. 10 μL was pipetted to the center of each well in a 48-well tissue culture-treated plate (Corning, 353078) and allowed to polymerize for 1 hr in a humidified incubator (37°C, 5% CO_2_).

Collagen gels were overlayed with 200 µL of desired growth medium (Fig 2A). For SACs grown in plasma, undiluted pooled human plasma (Innovative Research, IPLALIH; 3.9532 mg/mL fibrinogen) was added onto collagen gels. SILAC RPMI 1640 (Gibco, A2494201) was supplemented with 0.426 mM lysine, 1.4 μM L-arginine, and 2.055 mM L-glutamine. RPMI was supplemented with glucose or fibrinogen to mimic healthy or obese-T2D levels. Healthy glucose = 5 mM. Hyperglycemic glucose = 15 mM. Healthy fibrinogen = 3 mg/mL. Obese fibrinogen = 6 mg/mL. Human AB serum (Valley Biomedical, HP1022HI) was supplemented to 5% total volume of RPMI media (summarized in Table S1). Collagen gel SACs were grown (37°C, 5% CO_2_, humidified) for indicated time points.

**Table 1.** *In vitro* SAC growth media. SILAC RPMI 1640 (Gibco, A2494201) containing 0.426 mM lysine, 1.4 μM L-arginine, and 2.055 mM L-glutamine was supplemented with glucose, fibrinogen, and serum to the described final concentrations for growing SACs in collagen gels. LG: low glucose; HG: high glucose; LG+S: low glucose with serum; HG+S: high glucose with serum; LGLF: low glucose, low fibrinogen; HGLF: high glucose, low fibrinogen; LGHF: low glucose, high fibrinogen; HGHF: high glucose, high fibrinogen.

| **Condition** | **Glucose (mM)** | **Fibrinogen (mg/mL)** | **Serum (%)** | **Growth structure** |
| --- | --- | --- | --- | --- |
| LG | 5 | 0 | 0 | cluster |
| HG | 15 | 0 | 0 | cluster |
| LG+S | 5 | 0 | 5 | cluster |
| HG+S | 15 | 0 | 5 | cluster |
| LGLF | 5 | 3 | 5 | SAC |
| HGLF | 15 | 3 | 5 | SAC |
| LGHF | 5 | 6 | 5 | SAC |
| HGHF | 15 | 6 | 5 | SAC |

SACs used for fluorescent imaging were grown in media with FITC-labeled human fibrinogen (Innovative Research, IHUFBGFITC). Plasma was supplemented with 1 mg/mL FITC-fibrinogen, resulting in ∼20% of total fibrinogen. In RPMI media, FITC-fibrinogen constituted 20% of supplemented fibrinogen.

To quantify SAC CFUs, gels were washed with sterile 1X PBS three times before SACs were counted. Collagen gels were gently lifted with sterile pipette tips and transferred to Lysing Matrix A tubes containing 1 mL of sterile 1X PBS. Samples were homogenized with a FastPrep-24 at 6.0 m/s for 40 s, serially diluted, and plated onto TSA. CFU counts were normalized to the numbers of SACs per well.

To determine the width of SAC and pseudocapsule for SILACplus grown SAC, images were measured in ImageJ. Measurements were completed in technical triplicate on three biological replicates per condition. QuPath (version 0.6.0) was used to analyze pseudocapsules surrounding plasma-grown SACs. SILACplus-grown SACs were excluded because incomplete pseudocapsules prevented accurate quantification of FITC- and DAPI-positive regions.

Pseudocapsule boundaries were manually annotated, and superpixels were generated within FITC and DAPI regions using the SLIC algorithm (σ = 5 pixels, spacing = 10 µm, 10 iterations, regularization = 0.25. Distance-to-boundary measurements were calculated for each superpixel, and pseudocapsule width was determined by summing the absolute distances to the FITC and DAPI border annotations. Surface area was determined by summing the total surface area of all superpixels within FITC and DAPI border annotations. Radius was determined by distance to FITC border.

### Minimum inhibitory concentration (MIC) assay

The MICs of gentamicin, vancomycin, and nafcillin with *S. aureus* strains USA300 and JAR06.01.31 were measured as previously described^23^. Overnight cultures were washed twice in PBS, resuspended to an OD_600_ of 0.7, and further diluted 1:100 in TSB. 100 µL of culture was mixed with 100 µL of 2-fold serially diluted antibiotics in TSB (final concentrations 0-256 μg/mL) in round-bottom 96-well plates. The plates were incubated with shaking at 37°C for 24 hr, and then bacterial growth was measured at OD_600_ with a BioTek ELx808 microplate reader. MICs were defined as the lowest concentration of antibiotics where turbidity was undetectable by OD_600_.

### SAC antibiotic tolerance

SACs grown for 24 hrs were washed with PBS three times. 200 µL of 100X MIC solutions of vancomycin, gentamicin, or nafcillin (Table S4) were added to SACs for 3 or 24 hrs. Antibiotic-treated collagen gels were rinsed thoroughly and processed for CFU enumeration, as described above. CFU data were normalized to PBS-treated control wells. For experiments with values over 100% the results were scaled to a 0-100 percent range.

(Antibiotic (Total log10 CFU / total # SAC) / PBS (Total log10 CFU / total # SAC)) * 100 = percent viable relative to PBS control.

### RPMI antibiotic diffusion assay

Low glucose, low fibrinogen RPMI media was prepared as previously described and 200 µL was aliquoted into 48 well plates and allowed to solidify for 24 hrs. The following day, solid media gels were treated with antibiotics at 100X the MIC for 3 hours. Media gels were washed with 1X PBS 3X, then lifted out of the wells and placed on a lawn of JAR06.01.31. Plates were incubated at 37°C overnight. PBS treated gels show no clearance. Antibiotic treated wells have zone of clearance, indicating retention of antibiotic in the solid media.

### *In vitro* SAC fixation and staining

To remove washed collagen gels containing SACs from 48-well plates, ∼300 µL of 6% agarose was added to each well and allowed to solidify for 10 min. The agarose plugs containing collagen gels and SACs were transferred to tissue embedding cassettes and submerged in 10% neutral buffered formalin (VIP-Fixative, Scigen 1060) for 24 hrs at room temperature. Briefly, samples were embedded in paraffin and thin-sectioned (5 µm thick). Slides were melted at 60°C overnight, deparaffinized with xylenes, and rehydrated. Samples submerged in 1:100 antigen retrieval solution (Vector Laboratories, H-3300) underwent heat induced epitope retrieval (HIER), as previously described^24^. Samples were washed twice in both diH_2_O and PBS for 5 min each, permeabilized with 0.5% TritonX100 in tris-buffered saline for 20 min at room temperature, then stained with 10 µg/mL DAPI (Invitrogen, 62248) in 1X PBS for 10 min. After two washes (PBS, 5 min), coverslips were mounted with ProLong™ Gold Antifade Mountant (Invitrogen, P36930).

### Microscopy

Brightfield images were acquired using an EVOS™ Digital Color Fluorescence microscope (Invitrogen). Confocal laser scanning microscopy (CLSM) was performed using a Leica Stellaris 5 microscope. CLSM images were acquired using a 63x oil immersion objective. Raw fluorescence images were uniformly contrast-adjusted in ImageJ (v2.14.0) using *Enhanced Contrast* with the following parameters: saturated pixels = 0.2% and “normalize” selected.

### Transmission Electron Microscopy

USA300 SACs were grown in SILACplus media for 24 hours. Mature SACs were fixed in osmium tetroxide, then dehydrated with a graded series of alcohols. Samples were infiltrated into liquid epoxy resin (Epon/Araldite or Spurr), embedded into molds, and polymerized at 70°C. Sample blocks were cut into 1.0-2.0 uM sections and stained with Toluidine Blue, then assessed with light microscopy to determine target area to be trimmed and thin sectioned (70 nm) for electron microscopy. Sections are then stained with uranyl acetate and lead citrate and examined under TEM.

### RNA extraction of *in vitro* SACs

Whole transcriptome analysis (RNA-Seq) was used to evaluate SAC responses to different media conditions (plasma, Table S1). Following 24 hr of growth, collagen gels containing SACs were washed twice with PBS. Only conditions without fibrinogen were stabilized with RNAprotect Bacteria Reagent (Qiagen). To ensure sufficient RNA quantities, eight technical replicates of collagen gels were pooled in Tris-EDTA buffer (12.5 mM Tris, 75 mM EDTA, pH 7.6) and acid phenol:chloroform (Ambion, AM9722) within Lysing Matrix B (MP Biomedicals, 1169110-CF) tubes. Samples were mechanically lysed with a FastPrep-24 at 6.0 m/s for 40 s, then phase separated by centrifugation (14,000 x g, 10 min). The aqueous phase was mixed with TRIzol, and RNA was purified using a Direct-zol RNA Miniprep kit (Zymo, R2052). Residual DNA was depleted using TURBO DNase (Invitrogen, AM2238).

### NGS Library Construction and Sequencing

Quality of the extracted total RNA was assessed on a Fragment Analyzer (Agilent Technologies, Inc.) and quantified using Qubit (ThermoFisher Scientific). RNAtag-Seq libraries were constructed as previously described^25^. Briefly, 200 ng of total RNA was fragmented, DNased treated, and dephosphorylated prior to the ligation of barcoded DNA adapters containing a 5’ phosphate and 3’ C3 spacer blocking group. The barcoded RNA fragments were pooled and cleaned using the RNA Clean and Concentrator-5 (ZymoBiomics). rRNA was depleted using the riboPOOL Pan-Bacterial depletion kit according to the manufacture’s specifications (siTOOLS BioTech). Depleted RNA pools were converted to cDNA using SMARTScribe reverse transcriptase (Takara Bio). The primers Ar2 and 3Tr3 were added to facilitate template switching^26^. Ar2 primes off the ligated DNA adapter and the 3Tr3 template switching oligo contains three guanosine bases complementary to the C-tail of the newly synthesized cDNA strand. 3Tr3 also contains a 5’ iso-deoxy CGC blocking group to prevent concatemer formation. The cDNA was amplified using Q5 high-fidelity polymerase (New England Biolabs) and unique dual indexing primers targeting regions of the ligated adapter and template switching oligo containing Illumina P5 and P7 sequences. Fragment size profiles and quantification of the libraries were measured using the Fragment Analyzer and Qubit, respectively. Library pools were sequenced on the Illumina NextSeq 2000 with paired-end reads of 63 and 54 cycles.

### RNA-Seq Analysis

Raw reads generated from the Illumina basecalls were demultiplexed at the pool-level using bcl2fastq version 2.19.0. Samples were further demultiplexed from their pool by the inline barcode (contained in read 1) using a custom R script. Quality filtering, inline barcode, and adapter removal were performed using Trim Galore version 0.6.2 with the following parameters: “--paired --fastqc --illumina --max_n 1 --clip_R1 9 –clip R2 6 --length 30”.

The processed reads were then mapped to *Staphylococcus aureus* USA300 FPR3757 (GCF_000013465.1) or *Staphylococcus aureus* JAR06.01.31 (GCF_051122515.1) genomes using bowtie2 version 2.3.5 with the following parameter “--very-sensitive”. Gene-level read quantification was derived using the subread-2.0.3 package (featureCounts) with a GTF annotation file and the following parameters: “-s 2 -t gene -g locus_tag”. Differential expression analysis was performed using DESeq2-1.28.1 with a p-value threshold of 0.05 within R version 4.0.2 (https://www.R-project.org/).

### RT-qPCR

RNA was reverse transcribed to cDNA using qScript cDNA synthesis reactions (Quantabio, 95047). qPCR was performed with PerfeCTa SYBR Green FastMix (95072, Quantabio) on a CFX Connect (Bio-Rad) with the following thermocycler program: 95°C for 3 minutes, 40 cycles of 95°C for 10 seconds followed by 62°C for 30 seconds, and a final melt curve from 65°C-95°C. qPCR primer sequences are listed in Table S2.

### Statistical analysis

Statistical analysis was performed with GraphPad Prism (v10.4.1) and R (v4.2.3). Statistical and multiple comparison tests are denoted in each figure legend.

## Results

### Increased *Staphylococcus aureus-*induced clotting in diabetic plasma compared to healthy plasma

*S. aureus* USA300 FPR3757^19^ hereafter USA300, and JAR06.01.31^20^ were grown in healthy and diabetic human plasma to determine the impact of the diabetic and obese environments on clot formation. Healthy patient samples had an A1C between 5.3 and 5.7 and a healthy body mass index (BMI, kg/m^2^) ^27^ of less than 24.9; diabetic individuals had elevated A1C between 7.0 - 7.9^14^ and overweight or obese BMI^27^ of > 25.5 or > 30.0, respectively (Table S1). A1C and BMI were significantly correlated (r = 0.7831, p = 0.0074) (Fig. 1A). Hereafter, patient samples will be referred to as “healthy” or “diabetic”.

**Figure 1.**
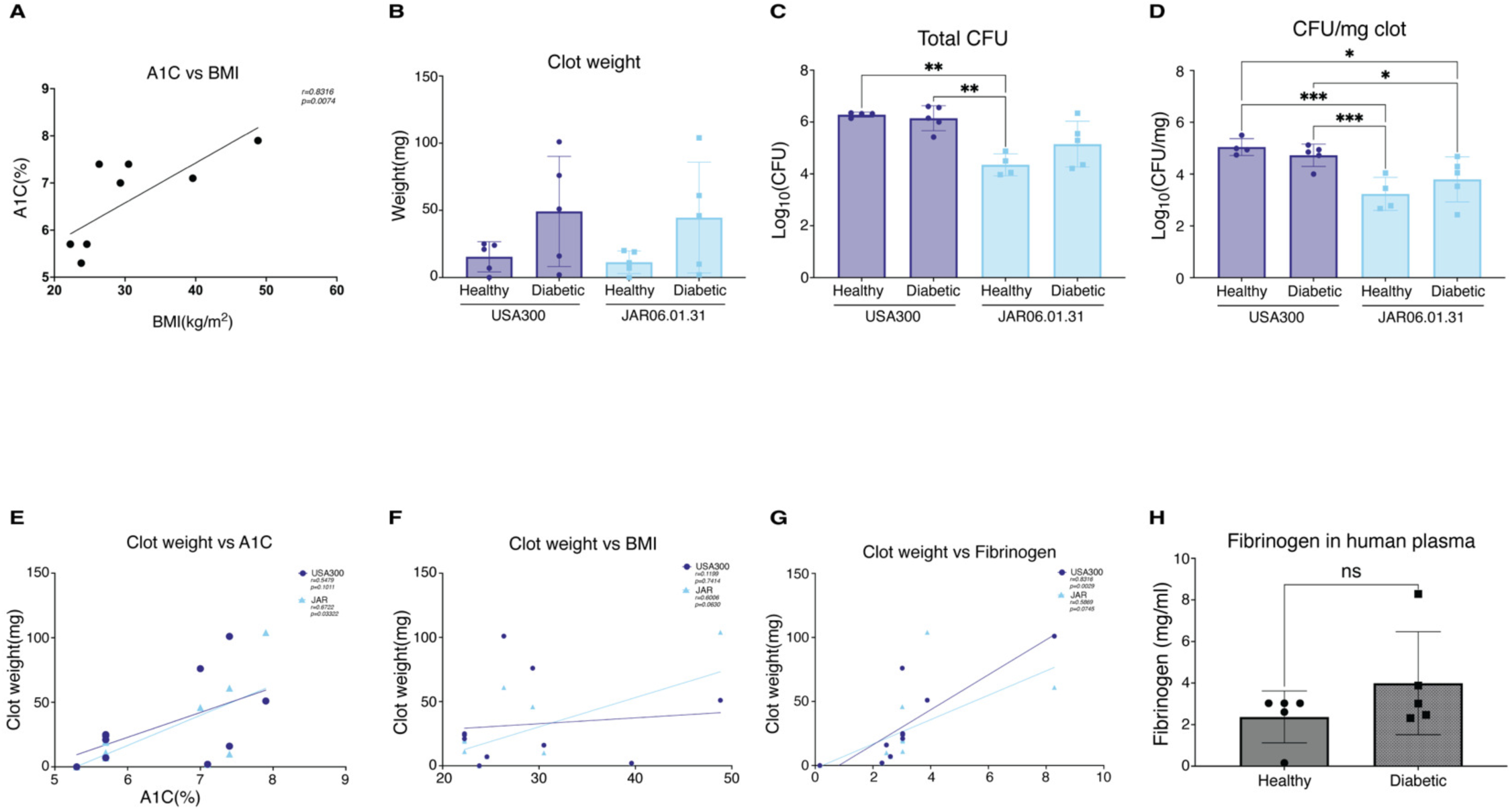
*S. aureus* forms larger clots in diabetic human plasma. **(A)** Correlation of patient A1C with BMI. Correlation was determined with Pearsons Correlation. N = 8. **(B-D)** Human plasma from healthy or diabetic patients was combined with approximately 1×10^8^ CFU of *S. aureus* USA300FPR3757 or JAR0601.31. Plasma and *S. aureus* were incubated at 37°C for 24 hours. Clot weight, total CFU per clot, and CFU/mg of clot were determined for each clot. N = 4-5. Significance determined with Two-way ANOVA with Tukey’s multiple comparison test. **(E-G)** Clot weight was correlated with A1C (%), BMI (kg/m^2^), and plasma fibrinogen concentrations (mg/mL). Correlation was determined with Pearsons Correlation. N = 16. **(H)** Fibrinogen concentrations of human plasma samples. Significance was determined with an Unpaired T-test. N = 4-5. All regression lines were determined with Simple linear regression. (* for p < 0.05, ** for p < 0.01, and *** for p < 0.001)

We hypothesized that *S. aureus* forms larger clots in diabetic human plasma due to increased availability of fibrinogen and host matrix factors (HMFs)^11,12^. Both strains formed heavier clots in diabetic plasma, though differences were not statistically significant (Fig. 1B). Total CFU did not significantly differ between healthy and diabetic plasma (Fig. 1C), consistent with growth in high-glucose media (Fig. S1), although JAR06.01.31 showed a trend toward higher CFUs in diabetic plasma compared to healthy plasma. USA300 formed clots with significantly higher CFUs than JAR06.01.31, suggesting enhanced adaptation to the hyperglycemic host environment (Fig. 1C). When CFUs were normalized to clot weight, less of the clot mass in JAR06.01.31 clots were due to bacterial load, suggesting a greater contribution of HMFs to clot weight (Fig. 1D). Similar trends were observed in mouse plasma (Fig. S2).

We assessed the correlation between clot weight and A1C, BMI, and fibrinogen concentration (mg/mL) in human plasma (Fig. 1E-G). JAR06.01.31 clot weight positively correlated with A1C, suggesting this strain may adapt to elevated glucose during clot formation. USA300 showed no significant correlation with A1C (Fig. 1E). Neither USA300 nor JAR06.01.31 clot weight correlated significantly with BMI (Fig. 1F). USA300 clot weight showed a strong positive correlation with fibrinogen concentration while JAR06.01.31 did not (Fig. 1G), suggesting USA300 may better adapt to elevated fibrinogen levels. Importantly, fibrinogen concentrations were not significantly different between healthy and diabetic plasma samples (Fig. 1H). Although all patients had an elevated A1C and uncontrolled diabetes, not all patients were obese. It has been shown that fibrinogen is associated with the degree of obesity, thus examining samples from patients with BMI > 30 may elucidate differences in fibrinogen levels^28^.

In summary, both strains formed larger clots in diabetic plasma. Clot weight of USA300 correlated with fibrinogen concentrations, indicating enhanced recruitment and utilization of fibrin. In contrast, JAR06.01.31 clot weight correlated with A1C, suggesting adaptation to elevated glucose in diabetic conditions. Since fibrinogen concentrations were similar between healthy and diabetic plasma, these findings indicate that strain-dependent differences in how *S. aureus* exploits host factors^29,30^, rather than fibrinogen abundance alone, drive enhanced clot formation in diabetic plasma.

### JAR06.01.31 SACs form a thicker fibrin pseudocapsule than USA300 SACs in human plasma

To assess staphylococcal abscess community (SAC) development *in vitro*, SACs were grown in 3D collagen gel matrices overlaid with pooled healthy human plasma or cell culture media (Fig. 2A, Table 1)^18^. Initial brightfield microscopy revealed differential SAC morphologies when comparing USA300 and JAR06.01.31 grown in healthy human plasma (Fig. 2B-C). The USA300 SAC fibrin pseudocapsules appeared as continuous defined spheres without a MAM^18^. In contrast, the JAR06.01.31 SAC fibrin pseudocapsules were significantly thicker and more fibrous with a MAM (Fig. 2B-C). The two strains yielded comparable CFUs per SAC (Fig. S3A), suggesting that the visual differences observed between SACs are due to differential fibrin or host matrix incorporation into the SAC pseudocapsule/MAM.

**Figure 2.**
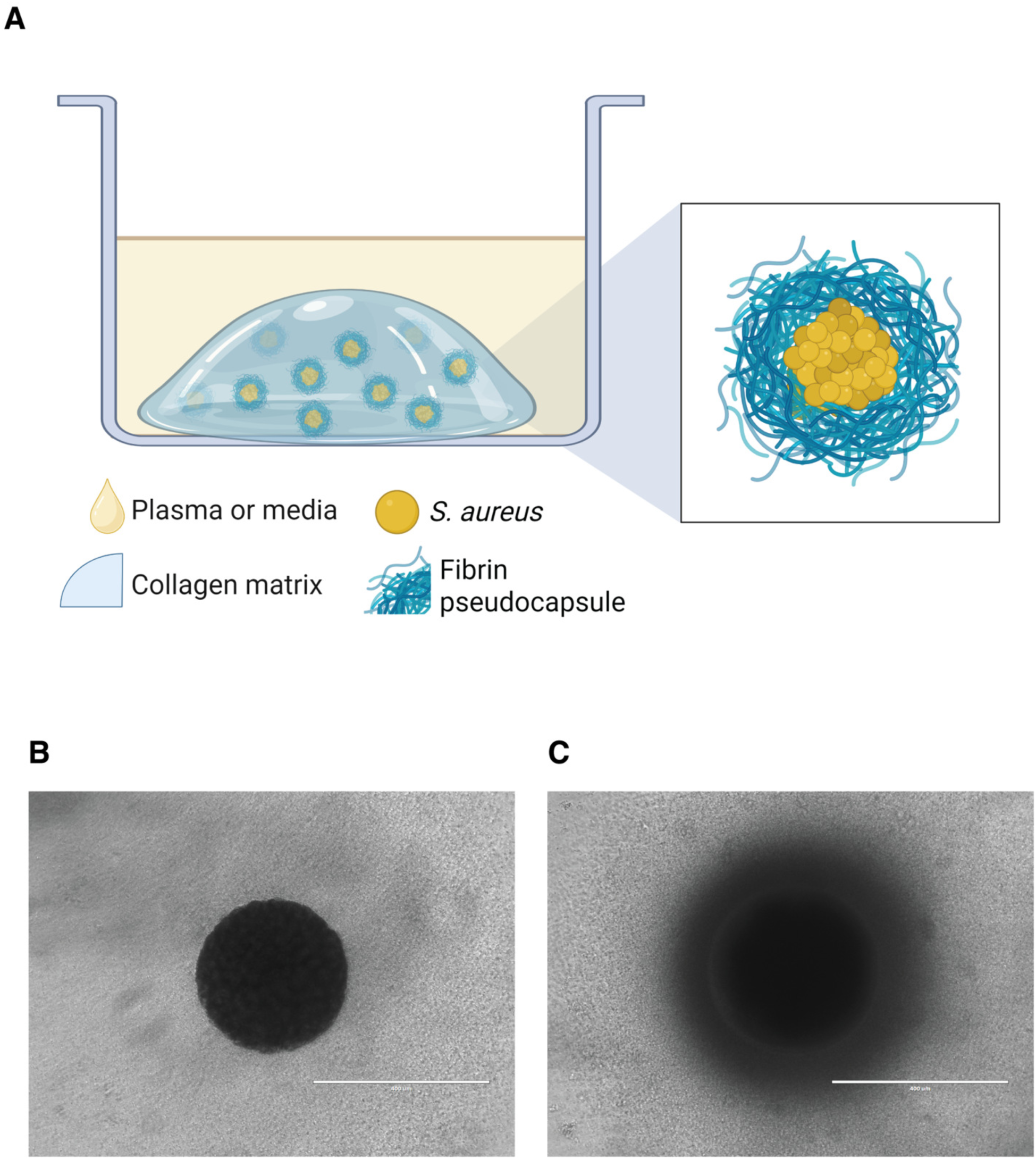
*In vitro* collagen gel assay for SAC growth. **(A)** Collagen gels mixed with approximately 20 CFU of *S. aureus* are incubated with pooled human plasma or SILACplus media (Table 1). As *S. aureus* grow within the collagen matrix, fibrinogen is recruited to form staphylococcal abscess communities (SACs). **(B, C)** Representative brightfield micrographs of *S. aureus* strain USA300FPR3757 and JAR06.01.31 SACs, respectively, grown with collagen gels in human plasma. Scale bars 400µm.

Based on our brightfield microscopy results and previous studies of *in vitro* SAC formation^31^, we hypothesized that USA300 and JAR06.01.31 employed unique mechanisms of fibrin pseudocapsule formation. To further assess these differences, we used confocal imaging of SACs grown in plasma supplemented with fluorescein isothiocyanate (FITC)-conjugated fibrinogen to visualize the fibrin pseudocapsule. Confocal images demonstrated clear differences between the size, pseudocapsule, and MAM width of USA300 and JAR06.01.31 SACs. The pseudocapsule boundary for JAR06.01.31 SACs was determined by density of the fibrin meshwork (Fig. S4). The area of the pseudocapsule was determined by FITC-positive signal within the pseudocapsule boundary (Fig. 3). USA300 SACs contained fibrin dispersed throughout the *S. aureus* core with a deposition of fibrin around the perimeter, forming the pseudocapsule (Fig. 3A-B). In contrast, JAR06.01.31 SACs lacked fibrin within the *S. aureus* core but had a greater deposition of fibrin around the perimeter, as well as a MAM (Fig. 3C, D). JAR06.01.31 SACs had a larger radius and pseudocapsule width than USA300 SACs (Fig. 3E, F). Furthermore, JAR06.01.31 SAC pseudocapsules had a greater contribution to the total radius of the SAC than pseudocapsules in USA300 SACs (Fig. 3G). Notably, the DAPI-positive area and total area of USA300 and JAR06.01.31 SACs are not statistically different, however, JAR06.01.31 has a greater FITC-positive area than USA300 SACs (Fig 3H-J). In summary, USA300 and JAR06.01.31 formed distinct SAC structures in plasma despite equivalent bacterial loads.

**Figure 3.**
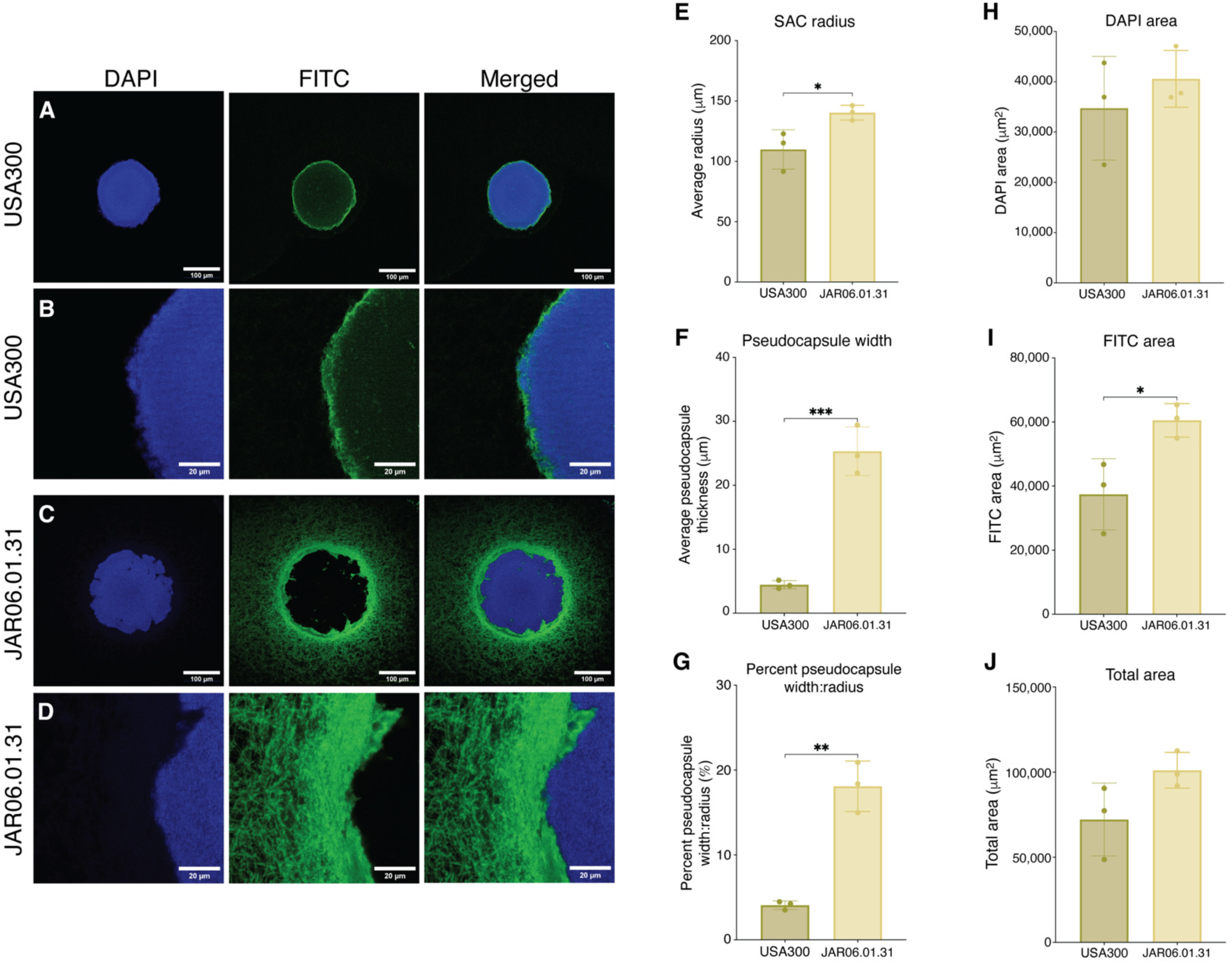
JAR06.01.31 SAC pseudocapsule is thicker than USA300 when grown in human plasma. Formalin fixed paraffin embedded sections of USA300FPR3757 and JAR06.01.31 SACs grown with FITC-conjugated fibrinogen in human plasma were stained with DAPI and confocal imaged. Images are representative of experiments in biological triplicate. **(A, B)** USA300. **(C, D)** JAR06.01.31. Scale bars **(A, C)** 100µm and **(B, D)** 20µm. **(E)** SAC radius determined as half of SAC diameter. **(F)** Pseudocapsule thickness determined as the width of FITC staining around SACs. **(G)** Percent of total radius width that is attributed to the pseudocapsule. **(H)** Area of DAPI+ area, **(I)** FITC+ area, and **(J)** FITC+ area normalized to DAPI+ area. Measurements were completed with QuPath. Pseudocapsule thickness and surface area per replicate were calculated from ∼300-600 superpixels and done in biological triplicate. Mean radius was calculated from annotations of DAPI positive and FITC positive regions in biological triplicate. Significance determined by Unpaired T-test (* for p<0.05, ** for p<0.01). All images were normalized in ImageJ.

### USA300 and JAR06.01.31 demonstrate alternative fibrinogen utilization strategies in simulated hyperglycemic and hyperfibrinogenemic conditions

Although not statistically significant, clot mass was greater in diabetic plasma than healthy plasma (Fig. 1) suggesting *S. aureus* adaptation to the obese/T2D environment. We hypothesized that *S. aureus* synergistically adapts to elevated glucose and fibrinogen availability to increase recruitment of fibrin to SAC structures. To explore this question, we grew *S. aureus* in collagen overlaid with SILACplus media (SILAC RPMI with lysine, arginine, glutamine) supplemented with 5% serum, glucose and/or fibrinogen at concentrations reflective of healthy (5 mM glucose, 3.0 mg/mL fibrinogen) and obese/T2D (15mM glucose, 6.0 mg/mL fibrinogen) physiological states^10,13^ (Table 1). *S. aureus* grown in low (healthy, LG) and high (obese/T2D, HG) glucose SILACplus media without serum and with serum (LG+S and HG+S) formed unstructured clusters of bacteria without a pseudocapsule (Fig. S5). Addition of serum and low (healthy) or high (obese/diabetic) levels of fibrinogen to low and high glucose SILACplus media (LGLF, LGHF, HGLF, and HGHF) was required for formation of a SAC structure with a surrounding pseudocapsule (Fig. 4), likely due to fibrin crosslinking initiated by serum prothrombin.

**Figure 4.**
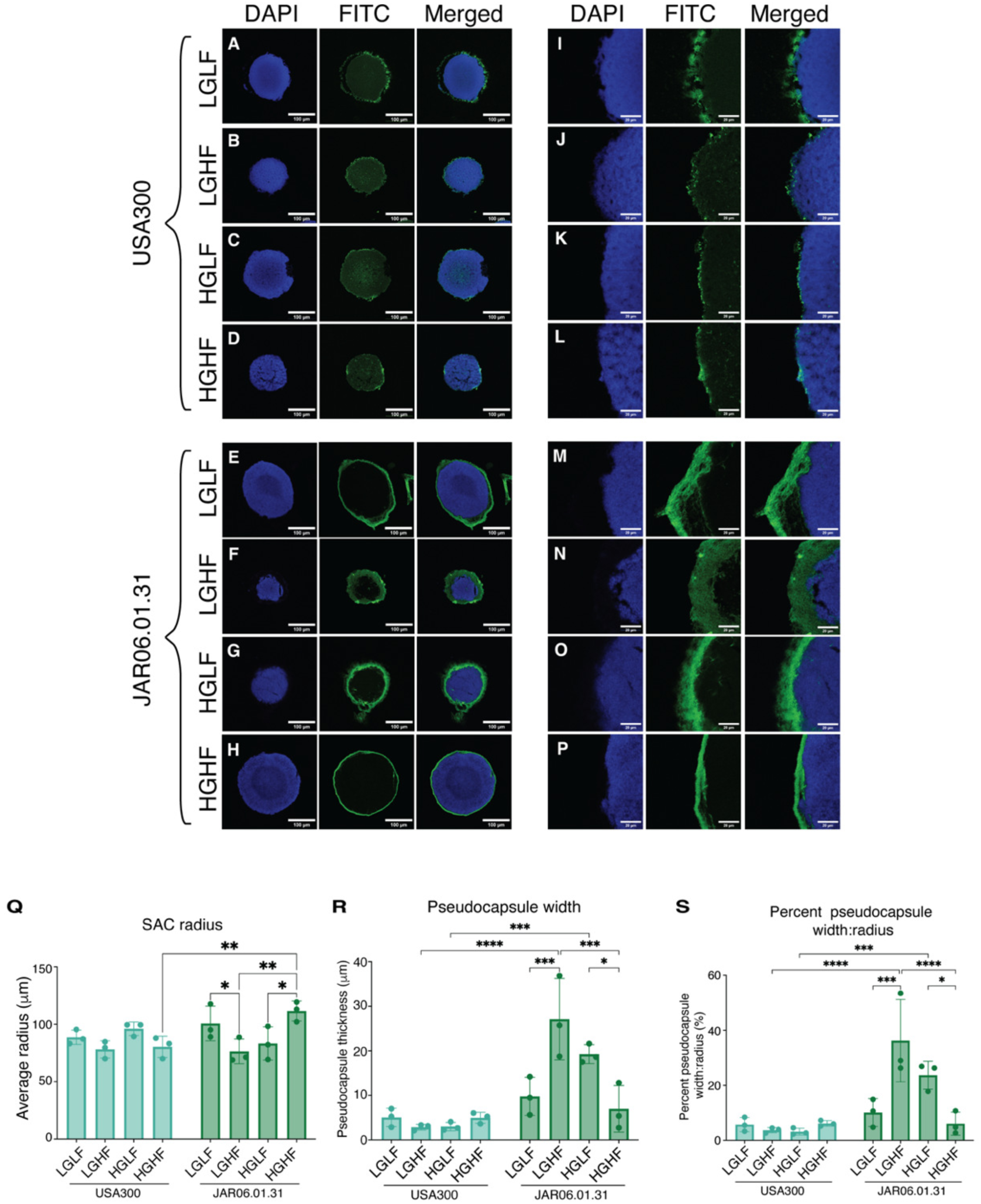
USA300 and JAR06.01.31 utilize fibrin differently in SAC formation. Formalin fixed paraffin embedded sections of USA300FPR3757 and JAR06.01.31 SACs grown with FITC-conjugated fibrinogen in SILACplus media supplemented with glucose (5 mM or 15 mM), fibrinogen (3 mg/mL or 6 mg/mL) and human serum (5%) (Table 1) were stained with DAPI and confocal imaged. Images are representative of experiments in biological triplicate. **(A-D, I-L)** USA300. **(E-H, M-P)** JAR06.01.31. Scale bars **(A-H)** 100µm and **(I-P)** 20µm. **(Q)** SAC radius determined as half of SAC diameter measured to the boundary of the pseudocapsule. **(R)** Pseudocapsule thickness determined as the width of FITC staining around SACs. Mean diameter and pseudocapsule thickness per replicate were calculated from six measurements on each SAC, done in technical and biological triplicate. **(S)** Pseudocapsule percentage of SAC radius. Significance was determined with Two-Way ANOVA with Tukey’s multiple comparison test (* for p < 0.05, ** for p < 0.01, and *** for p < 0.001). Healthy glucose = 5 mM. Hyperglycemic glucose = 15 mM. Healthy fibrinogen = 3 mg/mL. Obese fibrinogen = 6 mg/mL. **(A, I, E, M)** LGLF: low glucose, low fibrinogen; **(B, J, F, N)** LGHF: low glucose, high fibrinogen. **(C, K, G, O)** HGLF: high glucose, low fibrinogen; **(D, L, H, P)** HGHF: high glucose, high fibrinogen. Some images were rotated in image J for consistency. All images were normalized in ImageJ.

In all SILACplus media conditions, USA300 had significantly higher CFUs in each cluster or SAC than JAR06.01.31 (Fig. S3B), consistent with findings from stationary clotting assays (Fig. 1C). This suggests that JAR06.01.31 is at a growth disadvantage, with distinct nutrient utilization strategies under nutrient limited SILACplus conditions. Both USA300 and JAR06.01.31 grown in LG+S and HG+S SILACplus had greater CFUs in each cluster than in serum-free SILACplus (Fig. S3B). Elevated CFU counts may reflect the enhanced availability of alternative carbon sources, which *S. aureus* can metabolize to support growth and influence infection outcomes^32–36^. Similar to the differences in CFUs, there were more clusters of USA300 than JAR06.01.31 in each collagen gel in HG and HG+S SILACplus media (Fig. S3C). This suggests that elevated glucose concentrations enhance *in vitro* formation of USA300 clusters. However, this potential advantage is lost upon the addition of fibrinogen (Fig. S3C).

To further investigate if *S. aureus* synergistically adapts to elevated glucose and fibrinogen availability, confocal microscopy was used to examine SACs grown in SILACplus media supplemented with FITC-fibrinogen. Similar to observations in human plasma (Fig. 3), USA300 and JAR06.01.31 SACs formed in SILACplus media demonstrated distinct fibrinogen utilization strategies (Fig. 4). USA300 SACs had fibrin throughout the *S. aureus* core with discontinuous fibrin deposition around the perimeter, suggesting that USA300 recruits and integrates fibrinogen within the *S. aureus* core during SAC formation (Fig. 4A-D, I-L, Fig. S6). In contrast, JAR06.0.31 SACs lacked an internal gradient of fibrin and incorporated fibrin exclusively in the pseudocapsule surrounding the *S. aureus* core (Fig. 4E-H, M-P). Notably, JAR06.01.31 SACs grown in SILACplus conditions do not form a MAM (Fig. 4M-P). The fragmented nature of the USA300 pseudocapsule and lack of a MAM surrounding JAR06.01.31 SACs may suggest a role for host factors present in plasma but absent in SILACplus.

We next determined if SAC size and pseudocapsule thickness depends on fibrinogen availability (Fig. 4Q-S). USA300 SAC radius was smaller in high fibrinogen conditions (LGHF and HGHF) relative to low fibrinogen conditions (LGLF and HGLF), although the difference is not statistically significant (Fig. 4Q). Pseudocapsule width and the relative contribution of pseudocapsule to the total radius was not statistically different for either fibrinogen concentration (Fig. 4R, S), suggesting a threshold to fibrinogen incorporation. A possible maximum saturation of fibrinogen remains to be experimentally tested.

JAR06.01.31 SAC radius was significantly smaller in LGHF and HGLF compared to LGLF and HGHF (Fig. 4Q). JAR06.01.31 HGHF SAC radius was significantly greater than USA300 HGHF SAC radius (Fig. 4Q). JAR06.01.31 SACs demonstrated more variable pseudocapsule characteristics across SILACplus conditions (Fig. 4E-H, M-P). Despite smaller SAC radius in LGHF vs LFLF SACs, pseudocapsule width was greatest in these conditions (Fig. 4R), and a larger proportion of the pseudocapsule was attributed to the total SAC radius (Fig 4S). We found the opposite effect in high glucose conditions, where HGHF SACs had smaller pseudocapsule width (Fig. 4R) and smaller contribution of the pseudocapsule to the total radius (Fig. 4S) than HGLF SACs.

Together, this suggests that glucose availability modulates fibrinogen incorporation in a strain specific manner, and JAR06.01.31 is more successful adapting to and incorporating more fibrinogen in a low-glucose environment. In addition, SAC radius was not uniformly impacted by increases in glucose or fibrinogen (Fig. 4Q), highlighting the complex relationship between fibrinogen and nutrient availability in SAC formation. USA300 and JAR06.01.31 cluster size and uniformity was not significantly impacted by glucose concentration but was slightly affected by serum (Fig. S5).

Overall, USA300 SACs incorporated fibrinogen uniformly throughout the structure with a thin pseudocapsule; however, JAR06.01.31 SACs incorporated fibrin predominantly around the SAC perimeter with pseudocapsules thicker than those surrounding USA300 SACs. Furthermore, both strains responded differently to glucose and fibrinogen availability, which impacts SAC radius and pseudocapsule width. This may suggest differential regulation of genes involved in fibrin recruitment between the two strains in SILACplus media.

### Antibiotic-specific susceptibility modulated by SAC structure and metabolism

To compare antibiotic protection by the SAC pseudocapsule in healthy and obese/diabetic conditions, we exposed SACs and *S. aureus* clusters grown in hyperglycemic and hyperfibrinogenemic SILACplus media to 100X the MIC of gentamicin, vancomycin or nafcillin (Table S2). Planktonic USA300 and JAR06.01.31 cultures were susceptible to all antibiotics at 100X MIC (data not shown). We observed no significant differences in *S. aureus* USA300 survival after 3 hours of gentamicin treatment across all growth conditions, suggesting clusters and SACs conferred short-term protection (Fig. 5). Clusters grown in LG+S demonstrated a slight but not statistically significant increase in protection relative to LG alone, an effect not seen in HG+S. Notably, LGLF and LGHF provided significantly greater protection than HG+S (Fig. 5A). LGLF and LGHF conditions showed modestly increased survival compared to LG and LG+S conditions, though these differences did not reach statistical significance (Fig. 5A). A similar pattern was observed under high glucose conditions where HGLF and HGHF supported higher survival than HG or HG+S (Fig. 5A). These findings suggest that either concentration of fibrin is protective against gentamicin, and the metabolic state of the bacteria may impact survival. Gentamicin is an aminoglycoside that targets protein synthesis, thus lower metabolic activity in nutrient-poor low glucose conditions may increase tolerance and survival of *S. aureus*. However, this hypothesis remains to be investigated.

**Figure 5.**
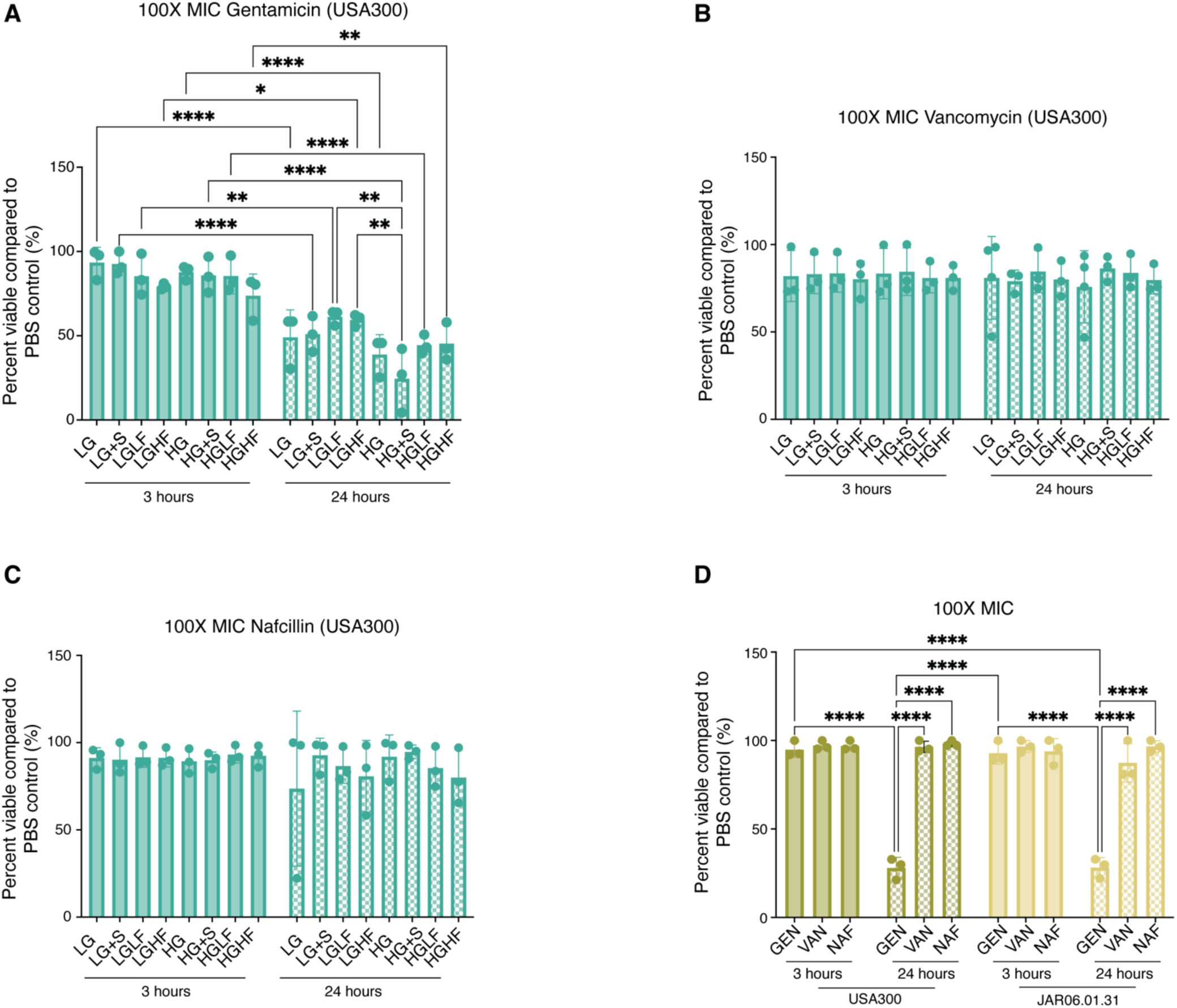
Fibrinogen alone may not provide protection to all antibiotics. *S. aureus* USA300FPR3757 and JAR06.01.31 were grown in **(A-C)** in SILACplus media supplemented with glucose (5 mM or 15 mM), fibrinogen (3 mg/mL or 6 mg/mL) and human serum (5%) (Table 1) or **(D)** pooled human plasma. SACs were grown as previously described then treated with antibiotics for 3 or 24 hours. Following antibiotic treatment, gels were washed thrice with sterile 1X PBS, homogenized, and serially diluted to enumerate CFU. GEN: gentamicin. NAF: nafcillin. VAN: vancomycin. Significance determined with Two-way ANOVA with Tukey’s multiple comparisons correction (* for p < 0.05, ** for p < 0.01, and *** for p < 0.001).

Vancomycin and nafcillin treatment produced no significant reduction in CFUs after either 3 or 24 hours across all tested conditions (Fig. 5B, C), indicating that clusters and SACs confer protection against vancomycin and nafcillin regardless of glucose or fibrinogen content. When grown in human plasma, USA300 SACs were resistant to treatment with gentamicin, vancomycin, and nafcillin for 3 hours (Fig. 5D). Protection was maintained after 24 hours of exposure to vancomycin and nafcillin, but viability was significantly reduced after 24 hours of treatment with gentamicin (Fig. 5D). This same trend was observed for JAR06.01.31 SACs grown in plasma. JAR06.01.031 SAC susceptibility to 400 µg/mL gentamicin builds upon previous reports that JAR06.01.31 SACs are resistant to 100 µg/mL gentamicin^31^.

In our initial experiments, SILACplus media supplemented with serum, glucose, and fibrinogen (LGLF, LGHF, HGLF, and HGHF; Table 1) solidified at room temperature, likely due to fibrin crosslinking initiated by serum prothrombin. When collagen gels were inoculated with USA300, the SILACplus media reliquefied, potentially caused by the production of staphylokinase (Sak), which activates plasminogen to degrade fibrin. This was verified by inoculation of the collagen matrix with a USA300Δ*sak* mutant where the SILACplus remained solid throughout growth (data not shown). Failure to reliquefy was also observed for gels inoculated with JAR06.01.31, which lacks *sak*. We determined that solidified SILACplus media retain antibiotics (Fig. S7A), thus JAR06.01.31 SACs in solid SILACplus media are exposed to antibiotics during homogenization prior to plating for enumeration of resistant colonies. As a result, an accurate determination of viability of JAR06.01.31 SACs exposed to antibiotics cannot be determined (Fig. S7B-D).

Together, these results demonstrate that the fibrin pseudocapsule and nutrient environment synergistically influence *S. aureus* survival during antibiotic exposure and highlight the need to consider both structure and metabolic state of clusters and SACs when evaluating antibiotic susceptibility.

### Gene expression of USA300 and JAR06.01.31 SACs in human plasma

We used bulk RNAseq to compare the gene expression of SACs grown in healthy human plasma to determine how each strain utilizes host matrix components during SAC formation (Table 1). Genes that were differentially expressed (|log2fold-change(log2FC)| > 3, p-adj < 0.05) in USA300 and JAR06.01.31 SACs are summarized in Fig. 6A.

**Figure 6.**
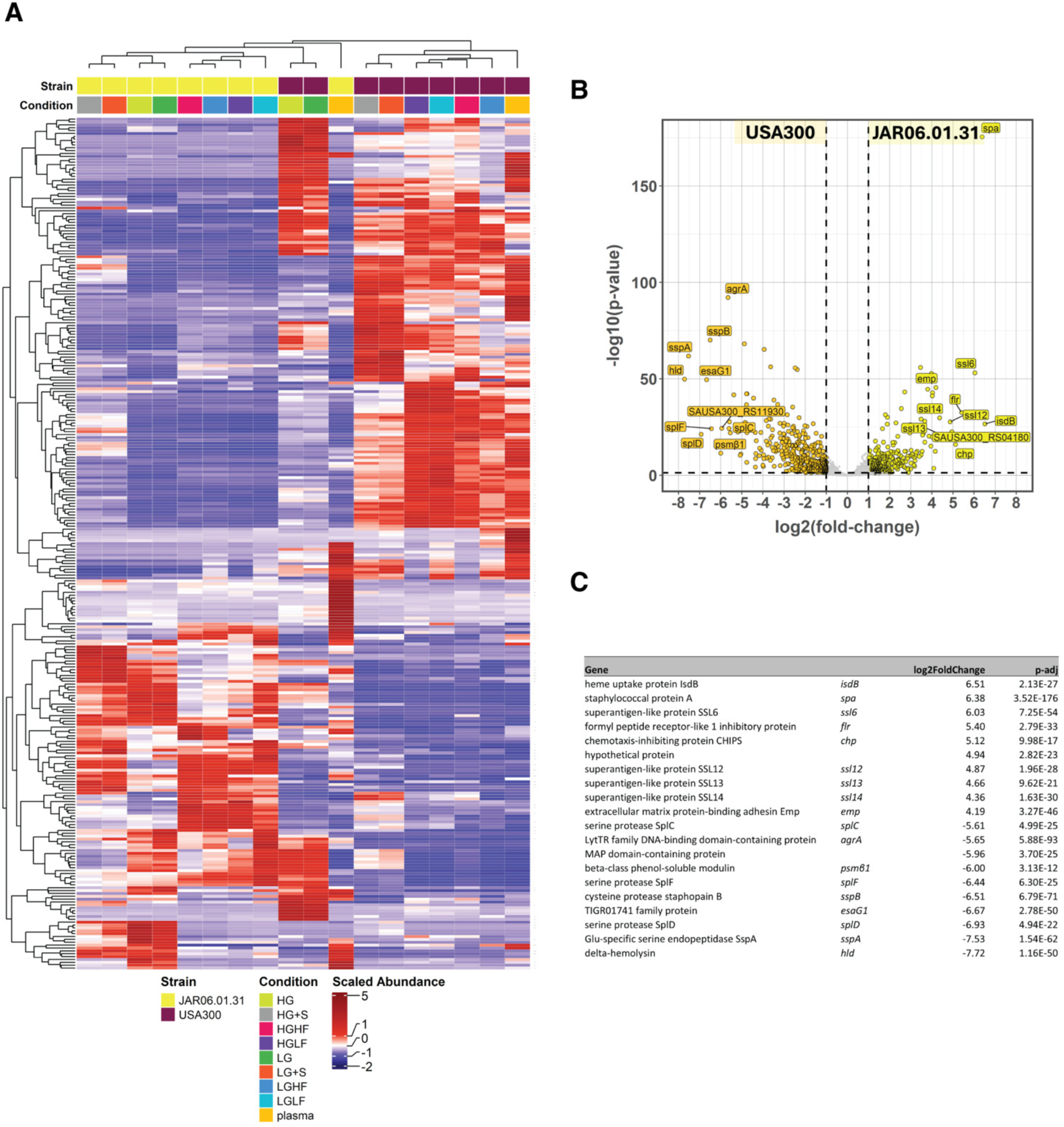
Gene expression of *S. aureus* JAR06.01.31 and USA300 within SACs varies with growth condition. **(A)** Heatmap represents significantly differentially expressed genes (|log2fold-change| > 3, p < 0.05) identified through RNAseq analysis of SACs grown in SILACplus media (Table 1). **(B)** Volcano plot represents differential gene expression of USA300 and JAR06.01.31 SACs grown in plasma, identified through DEseq2 differential expression analysis of RNAseq data. Significant (p-adj < 0.05) and |log2fold-change| > 1 values are shown in yellow (JAR06.01.31) or orange (USA300). The top 10 most differentially expressed genes according to magnitude are labeled and **(C)** listed with associated p-adj and log2fold-change values.

We first focused on gene expression of USA300 and JAR06.01.31 in human plasma to examine SACs in a healthy physiological host environment. As anticipated from distinct features of SAC formation that we observed in our brightfield and confocal images, the RNAseq data from USA300 and JAR06.01.31 SACs revealed 725 differentially expressed genes (DEGs) (|log2FC| > 1, p-adj < 0.05) (Table S3). Differential expression of the *S. aureus* accessory gene regulator (*agr*) global regulatory system and the functions controlled by *agr* accounted for the highest magnitude log2FC differences between the two strains (Fig. 6B). Regulatory activity of the Agr regulon is primarily via expression of RNAIII, which encodes delta-hemolysin (Hld), coordinates the upregulated expression of secreted exoenzymes and toxins, and downregulates expression of surface proteins^37–39^. The core components of the *agr* system include an autoinducing peptide (AgrD) which is transported out of the bacteria by a transmembrane endopeptidase (AgrB). AgrD is sensed by the cell wall-associated AgrC histidine kinase to activate AgrA, the response regulator for the Agr regulon, inducing expression of RNAIII. AgrA can also act independently of RNAIII to promote phenol-soluble modulin (PSM) transcription^40^.

In our results, expression of *agrA*, *hld*, and multiple α- and β-PSMs were differentially increased in USA300 compared to JAR06.01.31 when grown in healthy human plasma (Fig. 6B-C). Among the top 10 DEGs (log2FC) for USA300 were those encoding the *sspA, splD, sspB*, and *splF* proteases regulated by Agr^41,42^ (Fig. 6B-C). In comparison, JAR06.01.31 SACs grown in human plasma were enriched for transcripts that are repressed by *agr* activity, including surface protein-encoding genes (*isdB, spa, ssl6, ssl12, ssl13, ssl14, flr, chp*)^43–45^ (Fig. 6B-C). RT-qPCR for SAC expression of *RNAIII* and *spa* verified these RNAseq results (Fig. S8, Table S4).

Consistent with our observations, analysis of the JAR06.01.31 genome (GCF_051122515.1) revealed a four-nucleotide deletion resulting in a frameshift and premature stop codon within *agrC* (CP177129.1:1583984) (Fig. S9), which was predicted to render the two-component system histidine kinase ^46^ AgrC nonfunctional. High expression of exoproteins in JAR06.01.31 SACs may also be explained by differential expression of *sarS*^28^ (Table S3), which is a known activator of Spa^47,48^. Taken together, these data suggest that JAR06.01.31 *agr* is not functional and therefore relies on alternative regulatory systems such as the SarA family proteins to form and maintain SACs.

### Gene expression of USA300 and JAR06.01.31 SACs in the obese/T2D host environment

We next determined changes in gene expression of SACs grown in SILACplus media supplemented with serum, glucose, and fibrinogen to replicate the obese/T2D host environment (Table 1). Our initial analysis identified significant changes in gene expression depending on the glucose and fibrinogen concentrations (Fig. 6A). Comparisons of USA300 and JAR06.01.31 in all SILACplus conditions (Table 1) had similar top DEGs as in healthy human plasma (Table S5). To further characterize how SACs respond to varied concentrations of glucose and fibrinogen, gene expression data were clustered according to principal component analysis (PCA). For USA300 SACs, 83% of all variance in gene expression could be explained by two principal components (PC1 and PC2) (Fig. 7A).

**Figure 7.**
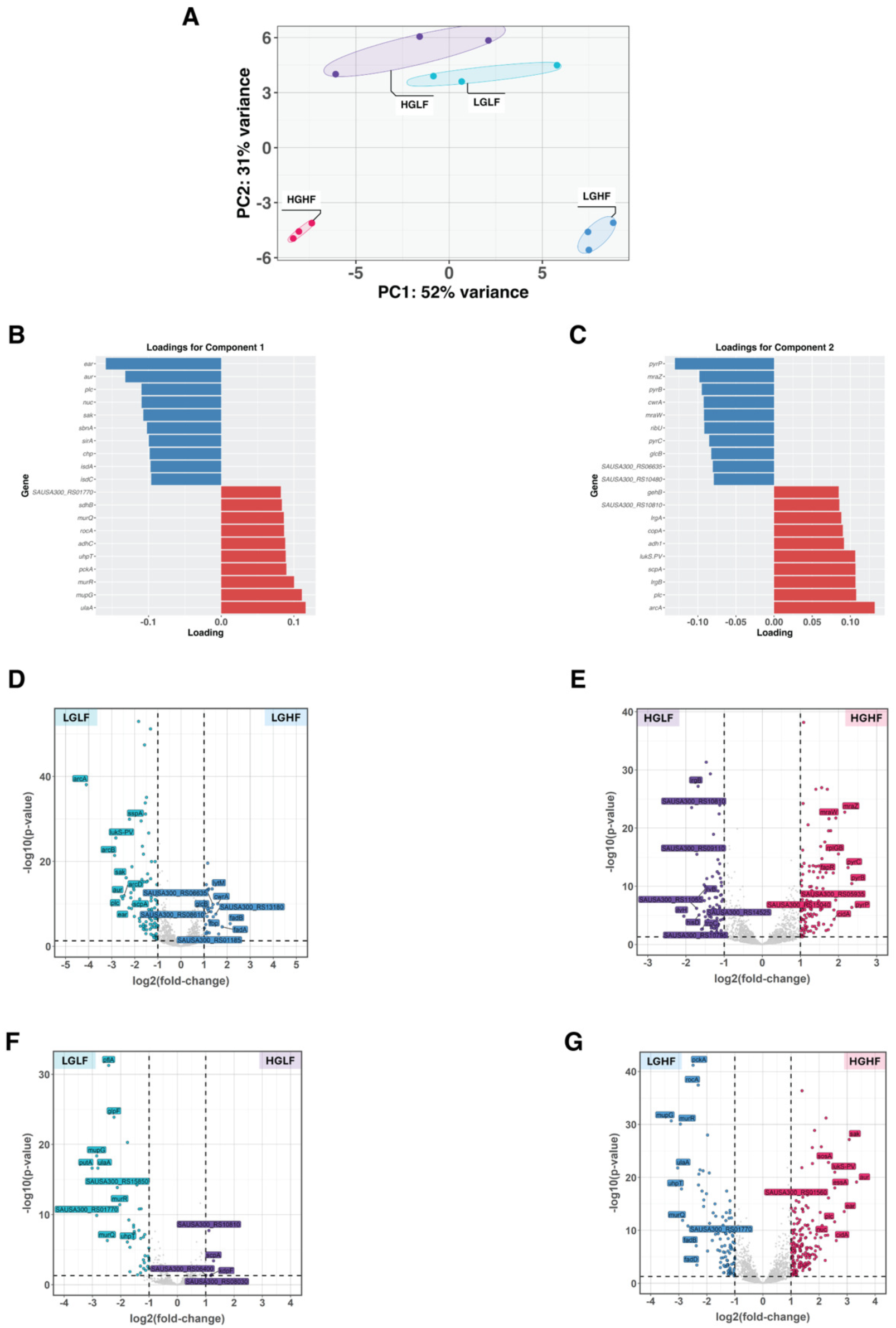
Fibrinogen and glucose concentrations contribute to differential gene expression in USA300 SACs. **(A)** Principal component analysis of the top 500 genes with the greatest variance across selected conditions in USA300. Bounding ellipses are drawn around individual conditions. **(B)** Top/bottom loading genes contributing to PC1. **(C)** Top/bottom loading genes contributing to PC2. Volcano plots represent differential gene expression of SACs grown in the indicated conditions, identified through RNAseq analysis. **(D)** Differentially expressed genes in USA300 SACs grown in LGHF compared to LGLF. **(E)** Differentially expressed genes in USA300 SACs grown in HGHF compared to HGLF. **(F)** Differentially expressed genes in USA300 SACs grown in HGLF compared to LGLF. **(G)** Differentially expressed genes in USA300 SACs grown in HGHF compared to LGHF. Significant (p < 0.05) and |log2fold-change| > 1 values are represented in color. The top 10 most differentially expressed genes according to magnitude are labeled.

USA300 SACs grown in media conditions differing in glucose concentration varied along PC1. PC1 loadings corresponding to LG are genes involved in metabolic and physiological processes like scavenging and metabolizing nutrients, especially non-canonical carbon sources, such as L-ascorbate (*ulaA*), MurNAc (*mupG, murR, murQ*) and host-derived glucose-6-phosphate (*uhpT*)^34,49–51^ (Fig. 7B). Upregulation of *pckA* and *sdhB* likely facilitate efficient ATP generation from alternative carbon sources^52,53^. Among these loadings, *ulaA, uhpT, pckA, murQ, and rocA* are regulated by catabolite control protein A (CcpA) and thus de-repressed during glucose scarcity^54–56^ (Fig. 7B).

Gene loadings associated with HG encode numerous tissue invasion and immune evasion factors, including *ear, aur, plc, nuc, sak*, and *chp* (Fig. 7B). Expression of these exotoxins represent a virulence repertoire driven by the Agr system that is acutely invasive and cytotoxic^57–60^. Additionally, HG was associated with genes involved in iron acquisition (*sbnA*, *sirA isdA, isdB*)^61^. Compared to LG conditions, USA300 SACs exhibited heightened virulence profiles in HG environments that would be predicted to exacerbate tissue invasion and bacterial survival *in vivo*.

In USA300 SACs, fibrinogen concentrations also impacted gene expression along PC2. Loadings associated with LF indicate a pathogenic state with expression of genes that contribute to immune evasion (*lukS-PV, scpA, plc, gehB)*^58,62^, autolysis (*lrgA, lrgB*)^63,64^, and other stress adaptations (*arcA, adh1, copA*)^65^ (Fig. 7C). These data reveal that SACs in LF conditions are stress-adapted, promoting tissue invasion, and expressing immunomodulatory factors that facilitate infection progression.

PC2 loadings associated with HF conditions include pyrimidine biosynthesis and salvage genes (*pyrP, pyrB, pyrC*)^66^ as well as genes that contribute to regulating the cell wall and division (*mraW, mraZ, cwrA*)^67,68^. Furthermore, *ribU* and *glcB* are associated with metabolism and nutrient acquisition which may facilitate survival in nutrient-poor or oxidative environments (Fig. 7C)^69^.

Comparing the virulence signatures of USA300 SACs grown in HF and LF, increased environmental fibrinogen concentrations alone do not promote enhanced virulence profiles. However, differential gene expression between paired media conditions shows that combinations of glucose and fibrinogen concentrations together impact SAC transcriptional profiles. The contribution of fibrinogen concentration, paired with low or high glucose, matches gene loadings identified for PC2 (Fig. 7C-E, Table S6) further supporting the conclusion that hyperfibrinogenemia alone is not a primary driver of *S. aureus* virulence in obese/T2D-like environments. Despite this, hyperfibrinogenemia paired with high glucose concentration results in a highly virulent transcriptional profile, which was not observed in LF conditions (Fig. 7F-G, Table S6). Genes encoding the Type VIIb secretion system, which contribute to pathogenesis of kidney abscesses *in vivo*^70^ were significantly differentially expressed in HGHF including *essA*, *essB, essC, esaA*,and *esxA* (Fig. 7G, Table S6). Taken together, these data illustrate that USA300 SACs acutely adapt gene expression according to environmental glucose and fibrinogen, with HGHF conditions resulting in a highly invasive and virulent transcriptional profile.

Furthermore, we examined how JAR06.01.31 SACs respond to hyperglycemic and hyperfibrinogenemic conditions via PCA. We found that JAR06.01.31 differentially responded to environmental glucose and fibrinogen to a lesser extent than USA300. LF and HF conditions were stratified along PC1 with minimal overlap. PC1 paralleled variations in fibrinogen concentrations and accounted for 44% of all variance (compared to 31% in USA300 PC2) (Fig. 8A). Notably, PC2 for JAR06.01.31 SAC gene expression did not align with changes resulting from glucose concentration. The gene loadings corresponding to HF for JAR06.01.31 SACs represent functions in *de novo* pyrimidine biosynthesis (*carA, carB, pyrB, pyrC, pyrD, pyrE, pyrF*) and stress adaptation (*ftnA, opuD2*)^71–73^ (Fig. 8B), indicating that SACs in HF are metabolically adapted to survive nutrient and osmotic stress.

**Figure 8.**
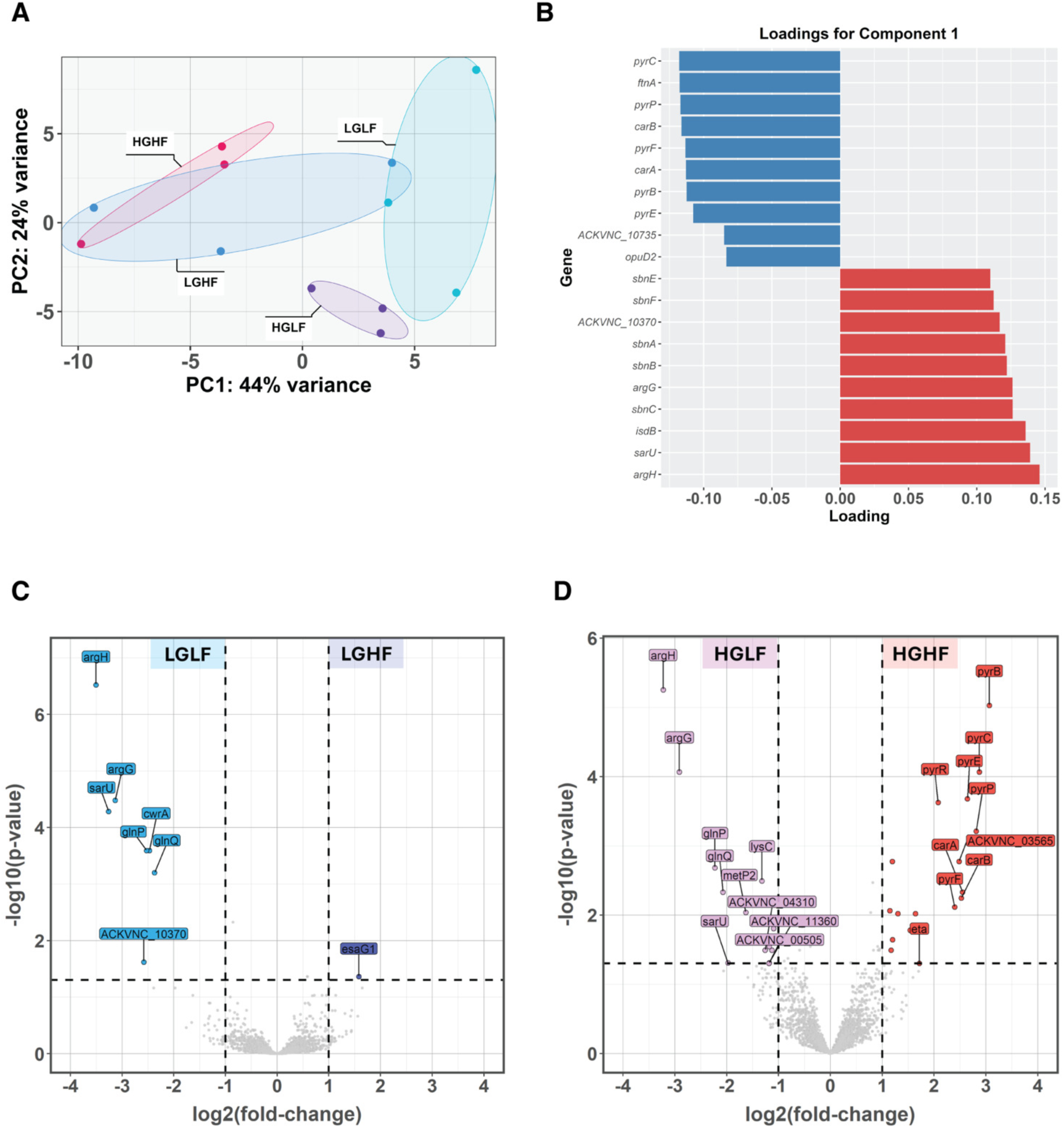
Fibrinogen and glucose concentrations contribute to differential gene expression in JAR06.01.31 SACs. **(A)** Principal component analysis of the top 500 genes with the greatest variance across selected conditions in JAR06.01.31. Bounding ellipses are drawn around individual conditions. **(B)** Top/bottom loading genes contributing to PC1. Volcano plots represent differential gene expression of SACs grown in the indicated conditions, identified through RNAseq analysis. **(C)** Differentially expressed genes in JAR06.01.31 SACs grown in LGHF compared to LGLF. **(D)** Differentially expressed genes in JAR06.01.31 SACs grown in HGHF compared to HGLF. Significant (p < 0.05) and |log2fold-change| > 1 values are represented in color. The top 10 most differentially expressed genes according to magnitude are labeled.

Gene loadings corresponding to LF indicate responses to nutritional immunity and a shift towards virulence priming. Genes that are associated with LF imply adaptation to limited arginine through arginine biosynthesis (*argG, argH*)^74^ as well as iron starvation through heme scavenging (*isdB*) and siderophore-mediated iron uptake (*sbnA, sbnB, sbnC, sbnE, sbnF*). Notably, *sarU* expression is reported to enhance virulence determinants in *S. aureus* by partially bypassing Agr to promote RNAIII^75,76^.

Although glucose concentration did not align with a component on PCA, the contribution of fibrinogen to JAR06.01.31 SAC differential gene expression was altered by LG or HG. Consistent with gene loadings for PC1 (Fig. 8B), SACs in LF conditions differentially expressed amino acid synthesis and uptake genes along with *sarU* in both LG and HG (Fig. 8C-D, Table S7). DEGs associated with HF are almost exclusively within HG conditions and consist of pyrimidine biosynthetic genes (Fig. 8D, Table S7). HGHF nucleotide metabolism may indicate a replicative phenotype, as opposed to a more persistent survival phenotype in LF conditions. Taken together, these data demonstrate that JAR06.01.31 transcriptional responses are primarily driven by fibrinogen rather than glucose; low fibrinogen was associated with virulence priming and iron and arginine acquisition, while high fibrinogen was associated with nucleotide metabolism and stress adaptation.

## Discussion

The hyperglycemic and hyperfibrinogenemic environment of obesity-related type 2 diabetes (obesity/T2D) contributes to a greater risk of *S. aureus* invasion and chronic infection. Addressing how *S. aureus* adapts to obese/T2D is crucial to understanding the pathogen-driven processes that enhance abscess formation and dissemination in this host environment. Our results shed light on this question by demonstrating that two clinically relevant *S. aureus* isolates differentially utilize and recruit host fibrin to form *in vitro* abscesses, expanding the current knowledge on adaptation of *S. aureus* to specific host states that are known to enhance virulence.

Our initial investigations in plasma clotting, an initial step for SAC formation, revealed that both USA300 and JAR06.01.31 formed larger clots in diabetic plasma compared to healthy plasma. For USA300, clot weight correlated with fibrinogen concentrations, indicating enhanced recruitment and utilization of host extracellular matrix components found in plasma, including fibrinogen. *S. aureus* JAR06.01.31 clot weight correlated with A1C, and not fibrinogen, suggesting adaptation to elevated glucose in diabetic conditions (Fig. 1). Since fibrinogen concentrations were not elevated in diabetic plasma, these findings indicate that other host factors^29,30^ beyond fibrinogen alone drive enhanced clot formation in diabetic plasma. In addition to increased fibrinogen and thrombin generation potential in obesity/T2D^77^, *S. aureus* maintains a vast repertoire of adhesins that bind components of the host extracellular matrix, including collagen, elastin, fibronectin, and laminin^30^, potentially explaining increased clot formation in JAR06.01.31 diabetic plasma clots despite no correlation between clot weight and fibrinogen concentration.

Previous studies have shown that SAC structure included a fibrin pseudocapsule and a fibrin microcolony-associated meshwork (MAM)^18^. It remains unclear how distinct MAM structure is from the pseudocapsule and if it has a role in *in vivo* SAC formation or protection from immune infiltrate^78^. For this reason, we annotated fibrin deposition at the SAC periphery as the pseudocapsule structure (Fig. 3, S4). USA300 formed smaller SACs with thinner pseudocapsules, incorporated fibrinogen throughout the abscess structure with increased density toward the perimeter of the SAC and lacked a MAM structure. In contrast, JAR06.01.31 formed larger SACs with thicker pseudocapsules, incorporated the majority of fibrin around the perimeter of the SAC, and formed a MAM, demonstrating unique fibrin utilization strategies consistent with previous observations of strain-to-strain differences in SAC architecture^18,31^. The increased expression of surface protein-encoding genes (*isdB, spa, ssl6, ssl12, ssl13, ssl14, flr, chp*) in JAR06.01.31 SACs grown in human plasma potentially provide mechanisms for increased recruitment of host extracellular matrix components^79,80^.

In plasma and in SILACplus, USA300 incorporates fibrin throughout the SAC structure whereas JAR06.01.31 exclusively incorporates fibrin around the perimeter of the SAC. Similar JAR06.01.31 pseudocapsule morphology, including thickness and MAM structure, has previously been observed^31^. We found that USA300 SACs grown in human plasma have a more consistent pseudocapsule, without gaps, than USA300 SACs grown in SILACplus media (Fig. 3, 4). This is likely due to other host factors in human plasma including collagen, elastin, and fibronectin, that may enhance pseudocapsule formation^30^. In support of this, we found that USA300 SACs grown in SILACplus conditions do not consistently form an intact pseudocapsule (Fig. 4). This suggests that other host matrix components are required for SAC pseudocapsule formation in USA300. In human plasma, JAR06.01.31 forms a thick pseudocapsule with a MAM. Interestingly, JAR06.01.31 forms a thick uninterrupted pseudocapsule in SILACplus media, but lacks the MAM seen in plasma SACs. This suggests differences in pseudocapsule formation strategies between the two strains. We also observed differences in cluster radius based on serum (Fig. S5) and SAC radius based on glucose and fibrinogen availability (Fig. 4).

Antibiotic susceptibility testing revealed plasma and SILACplus SACs are protected from both nafcillin and vancomycin. Previous work has shown that antibiotics are localized around the fibrin pseudocapsule of *S. aureus* SACs^31^. We hypothesize this protection stems from the physical exclusion by size and charge of antibiotics^81^ around the fibrin pseudocapsule. The net negative charge of the pseudocapsule^82^ may prevent infiltration by smaller negatively charged antibiotics like nafcillin. Size may be the primary barrier for larger, positively charged antibiotics, such as vancomycin. In contrast, positively charged and small antibiotics like gentamicin may be able to infiltrate the pseudocapsule. Alternatively, in the nutrient-limited SAC environment, *S. aureus* may enter a quiescent metabolic state, which has been shown to promote antibiotic tolerance ^83–85^ and impair immune responses to infection^85,86^. We speculate that *S. aureus* clusters retained viability throughout antibiotic exposure despite lacking a pseudocapsule by adopting a dormant-like state, as observed contributing to biofilm-mediated antimicrobial resistance^87^. Similarly, extracellular DNA or other secreted products may contribute to protective matrices encompassing *S. aureus* clusters. It has also been shown that human serum alone can trigger antibiotic tolerance in *S. aureus* through increased peptidoglycan and cardiolipin accumulation^88^; however, this hypothesis remains to be elucidated in this model. Overall, these data indicate that the three-dimensional structure of the clusters and associated metabolic shifts confer protection, even in the absence of serum and fibrinogen. Interestingly, SILACplus SAC susceptibility to gentamicin is not greatly impacted by glucose or fibrinogen concentrations. Although fibrinogen availability does not seem to directly impact antibiotic protection, a better understanding of how the pseudocapsule protects *S. aureus* against antibiotics is important to inform treatment strategies.

RNA-seq analysis revealed that SAC transcriptional profiles are shaped by both strain background and environmental glucose and fibrinogen concentration. Under high glucose, USA300 SACs showed elevated *agr* activity^37–39^ with upregulation of exotoxins, proteases, and virulence genes, while low glucose promoted expression of metabolic genes for alternative carbon sources. Fibrinogen availability further stratified gene expression with low fibrinogen associated with stress adaptation and immune evasion, and high fibrinogen linked to nucleotide biosynthesis, cell wall regulation, and survival in nutrient-poor or oxidative environments. This aligns with *S. aureus* adapting to the chronically inflamed T2D host, where obesity exacerbates chronic oxidative stress^89^. Notably, the greatest difference in gene expression between low glucose samples and high glucose samples occur only when fibrinogen is also high (HGHF vs. LGHF, Fig. 7G). This suggests that within the hypercoagulable environment of obese/T2D hosts with hyperglycemia, controlling blood sugar alone might help prevent excessive exotoxin production by *S. aureus*.

In contrast, JAR06.01.31 gene expression is primarily driven by fibrinogen. Minimal response to glucose concentration is consistent with host-adapted Agr dysfunction reported in persistent infections^90,91^. Mutation in *agrC* specifically, as we observed in JAR06.01.31, is reported to reduce virulence while promoting persistence functions^92,93^. Here, low fibrinogen promoted virulence priming through *sarU* and iron and arginine acquisition, while high fibrinogen supported nucleotide metabolism and stress adaptation. JAR06.01.31 lacks Sak, the fibrin-specific plasminogen activator that is positively regulated by Agr and secreted by *S. aureus* strains like USA300^19,94,95^. Sak-mediated fibrinolysis has been proposed to control SAC dissemination, which, when considering the lack of Sak in JAR06.01.31, may partially explain why increased environmental fibrinogen leads to stress-adapted persistence phenotypes for JAR06.01.31 SACs^18,31,96^. In *S. aureus* biofilms, both the Agr system and Sak have roles in regulating biofilm formation and dispersal by mediating fibrinolysis and other proteolytic degradation within the biofilm matrix^97,98^. Staphylococci lacking *agr* are reported to form thicker biofilms, since cells cannot detach from mature biofilms, and the pathogenicity of these strains varies^99–101^.

Together, these observations support a model in which *S. aureus* exploits both the hyperglycemic and procoagulant hyperfibrinogenemic environment of obese/T2D hosts. Hyperglycemia heightens expression of *S. aureus* virulence determinants, while hyperfibrinogenemia and enhanced clotting support of abscess architecture. This synergistic interaction likely contributes to the chronicity, severity, and recurrence of *S. aureus* infections in obese and diabetic populations and highlights the importance of considering the host metabolic state and coagulability status in management of *S. aureus* infections.

## Data Availability

The data discussed in this publication have been deposited in NCBI’s Gene Expression Omnibus^102^ and are accessible through GEO Series accession number GSE307544 (https://www.ncbi.nlm.nih.gov/geo/query/acc.cgi?acc=GSE307544).

## Contributions

E. B., L.M., T.B., and S.G. designed the research; B.R. collected the patient blood samples; E.B., L.M., and T.B. conducted the experiments; L.M. and A.L.G. conducted the mRNA sequencing and analysis; E.B. and L.M. performed the statistical analysis; E.B., L.M., T.B., A.L.G., and S.G. wrote the paper.

## Acknowledgements

This research was supported by NIH T32 GM135134, NIH T32 AI007285-37, and NIH R01 AR078414. The research reported in this publication also used resources and services provided by the University of Rochester Medical Center (URMC) Genomics Research Center (RRID:SCR_012359), the Histology, Biochemistry, and Molecular Imaging Core (NIH grants AR069655), the Center for Advanced Microscopy and Nanoscopy (RRID:SCR_023177), computational resources at the Center for Integrated Research Computing, and the Electron Microcopy Research Core (RRID:SCR_012366). Thank you to Karen de Mesy Bentley for electron microscopy support. Figure 2A “*In vitro* collagen gel assay for SAC growth” created in BioRender. Markle, L. (https://BioRender.com/4jxhosp) is licensed under CC BY 4.0.

